# The origin of fungi-culture in termites was associated with a shift to a mycolytic gut bacteria community

**DOI:** 10.1101/440172

**Authors:** Haofu Hu, Rafael Rodrigues da Costa, Bo Pilgaard, Morten Schiøtt, Lene Lange, Michael Poulsen

**Author notes:** contributed equally to this work (Haofu Hu, Rafael Rodrigues da Costa). Corresponding author Haofu Hu, Section for Ecology and Evolution, Department of Biology, University of Copenhagen, Universitetsparken 15, 2100 Copenhagen East, Denmark.

## Abstract

Termites forage on a range of substrates, and it has been suggested that diet shapes the composition and function of termite gut bacterial communities. Through comparative analyses of gut metagenomes in nine termite species with distinct diets, we characterise bacterial community compositions and identify biomass-degrading enzymes and the bacterial taxa that encode them. We find that fungus-growing termite guts are enriched in fungal cell wall-degrading and proteolytic enzymes, while wood-feeding termite gut communities are enriched for plant cell wall-degrading enzymes. Interestingly, wood-feeding termite gut bacteria code for abundant chitinolytic enzymes, suggesting that fungal biomass within the decaying wood likely contributes to gut bacteria or termite host nutrition. Across diets, the dominant biomass-degrading enzymes are predominantly coded for by the most abundant bacterial taxa, suggesting tight links between diet and gut community composition, with the most marked shift being the communities coding for the mycolytic capacity of the fungus-growing termite gut.

## Introduction

Termites are widespread in tropical, subtropical and warm temperate regions (Buczkowski and Bertelsmeir 2016) and form a diverse group of more than 3,000 described species in 281 genera and seven families (Eggleton 2000; 2001; Kambhampati and Eggleton 2000; Inward et al. 2007). They have major impacts on their environments (Buczkowski and Bertelsmeir, 2016), and this success has been attributed to their capacity to use nutritionally-imbalanced, recalcitrant food sources, allowing for colonization of otherwise inaccessible niches (Brune 2014). Different termites forage on distinct substrates, including soil, wood, dung, and fungus (Thongaram et al. 2005; Liu et al. 2013), decomposed through intricate interactions with complex gut microbial communities (Brune 2014; Brune and Dietrich 2015). In most termites, the main role of gut microbiota is believed to be the digestion of lignocellulose (Ni and Tokuda 2013; Watanabe and Tokuda 2010), but gut microbes also play key roles in nitrogen fixation (Breznak, 1982; Higashi et al. 1992; Sapountzis *et al.* 2016), microbial defence (e.g., Um et al. 2013) and immune regulation (Round and Mazmanian 2009; Viaud et al. 2014), of major importance for the evolutionary history of the symbioses.

Approximately 30 million years ago (MYA), the basal higher termite subfamily Macrotermitinae engaged in a mutualistic association with *Termitomyces* fungi (Aanen et al. 2002; Bourguignon et al. 2015) and shifted composition of the gut microbiota (Dietrich et al. 2014; Otani et al. 2014; 2016). *Termitomyces* decomposes plant material within external fungus gardens (combs) (Nobre et al. 2011; Poulsen et al. 2014), but the gut still remains central in the association, because plant substrate is macerated and mixed with asexual *Termitomyces* spores in a first gut passage prior to comb deposition (Leuthold et al. 1989). After *Termitomyces* breaks down the plant substrate, the termites ingest mature parts of the comb in a second gut passage (Leuthold et al. 1989), where gut microbes may contribute enzymes for final digestion of any remaining plant components (Poulsen et al. 2014). This division of labour is consistent with gut bacteria being of importance mainly when the comb material passes through the termite gut in a second passage cf.(Nobre et al. 2011; Poulsen et al. 2014; Poulsen, 2015), but recent work has suggested that partial lignin breakdown may also be accomplished during this first gut passage in *Odontotermes formosanus* (Li et al. 2017).

A set of microbes distinct from the gut microbiota of other termites persists in the fungus-growing termite guts, but limited work has examined the functional implications of this shift (Liu et al. 2013; Poulsen et al. 2014). It has been hypothesized that it was associated with the more protein-rich fungal diet (Dietrich et al. 2014) and/or to break down chitin and other fungal cell wall components (Hyodo et al. 2003 Liu et al. 2013; Poulsen et al. 2014). *Termitomyces* domestication exposed fungus-growing termite gut communities to large quantities of fungal cell wall glucans (composed of D-glucose monomers), chitin (glucosamine polymer), and glycoproteins (e.g., Reid and Bartnicki-Garcia 1976). Their breakdown requires a combination of carbohydrate-active enzymes (CAZymes; www.cazy.org; Cantarel et al. 2009; Lombard et al. 2014) and fungus-growing termite gut bacteria indeed encode glycoside hydrolase (GH) families of enzymes that may cleave chitin (GH18, GH19, and GH20), beta-glucan (GH55, GH81, and GH128), and alpha-mannan (GH38, GH76, GH92, GH99, and GH125) (Liu et al. 2013; Poulsen et al. 2014).

In nature, bacteria are the major chitin degraders and its hydrolysis has been correlated with bacterial abundances in e.g., soil communities (Kielak et al. 2013). In fungus-growing termites, Bacteroidetes and Firmicutes bacteria appear to be the main producers of CAZymes putatively producing mycolytic enzymes (Liu et al. 2013; Poulsen et al. 2014). These studies remained preliminary, however, because they were based on either an unassembled low-coverage metagenome (Liu et al. 2013) or had limited functional predictions (Poulsen et al. 2014). Here, we sequenced the gut metagenome of the fungus-growing termite *Odontotermes* sp. to elucidate its fungal and plant cell wall-degrading capacities at deeper functional levels (i.e., to EC numbers when possible), assigned putative enzyme functions to gut community members, and performed proteolytic enzyme analyses. To investigate the link between termite diet and gut community composition, we compared our findings to metagenomes from the fungus-growing termite *Macrotermes natalensis* (Poulsen et al. 2014) and seven non-fungus growing termite species feeding on plant material at different degrees of decomposition; the dung feeder: *Amitermes wheeleri* (He et al. 2013); two wood feeders: *Nasutitermes corniger* and*Microcerotermesparvus;* a litter feeder *Cornitermes* sp.; two humus feeders: *Termes hospes* and *Neocapritermes taracua;* and the soil feeder *Cubitermes ugandensis* (Rossmassler et al. 2015). We reveal that the shift in gut community composition after the origin of fungus farming was associated with a mycolytic microbiota, providing novel insights into digestion and the role of gut communities in the fungus-growing termite symbiosis.

## Results

### Taxonomic composition of fungus-growing termite gut microbiotas

We assigned bacterial taxonomies to metagenome contigs by searching for the closest matches of protein-coding genes on each contig against the NR database in NCBI and compared the relative abundance of the contigs in each group to assess the composition of termite gut microbiotas. *Ma. natalensis* and *Odontotermes* sp. were distinct from the seven other higher termite species with different diets (Figure 1A), corroborating previous work (Dietrich et al. 2014; Mikaelyan et al. 2015) that termites in the same feeding group are similar in gut microbiota composition (Figure 1B, Table S1). Firmicutes and Bacteroidetes dominated both *Odontotermes* sp. (24% and 25%, respectively) and *Ma. natalensis* (24% and 30%, respectively) (Figure 1A; Table S1), consistent with 16S rRNA amplicon surveys of these species (Otani et al. 2016) and metagenome analysis of *Odontotermesyunnanensis* from Southwest China (Liu et al. 2013). *Alistipes* and *Bacteroides* were the most abundant genera, representing respectively 5-8% and 2-4% total abundance, in sharp contrast to abundances in the other higher termites (on average 0.1% and 0.5%, respectively) (Table S1). *Treponema* (Spirochaetes) abundance was generally low in fungus growers and termites that feed on decaying plant material (3 and 6%, respectively) except for the litter feeder *Cornitermes* sp. (32%), while they were the most abundant taxon in wood feeders (42-45%) (Table S1).

**Figure 1.**
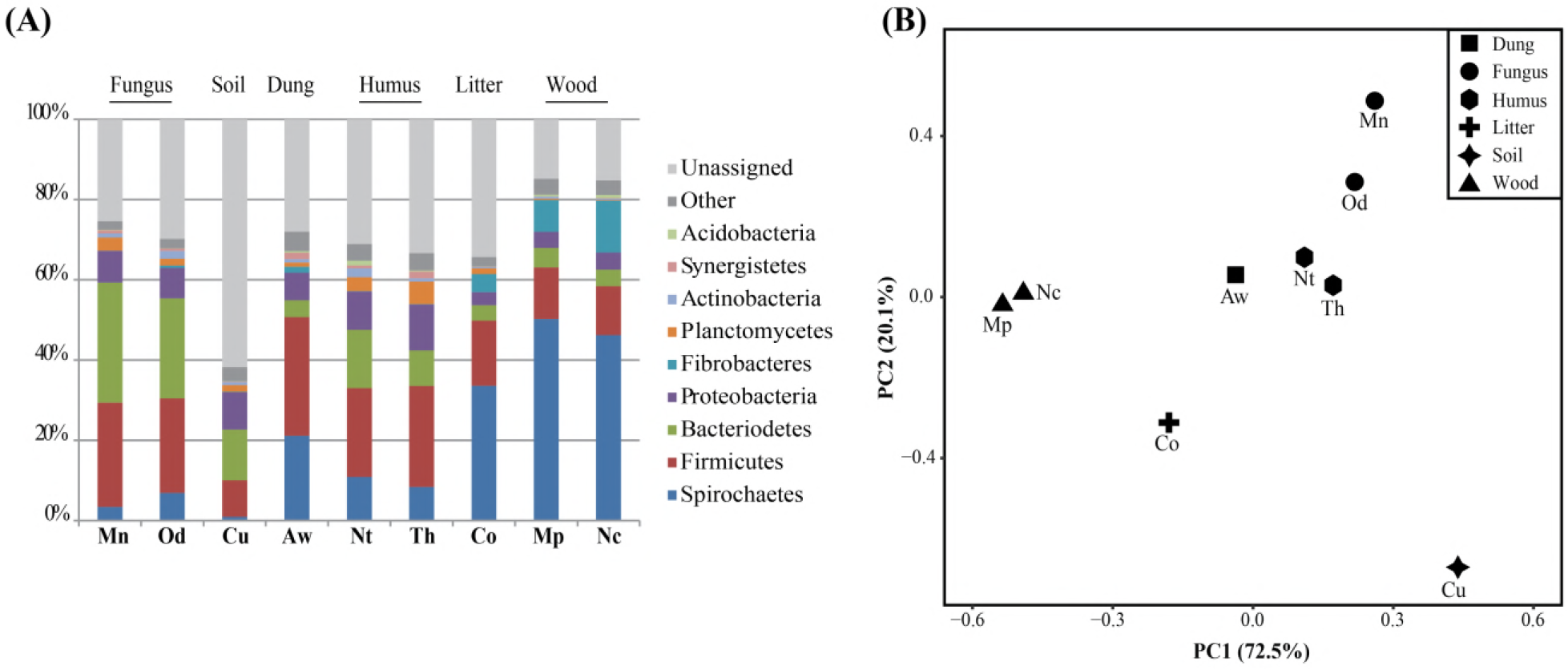
**(A)** Relative abundances of bacterial phyla each comprising more than 1% of the microbiota in the guts of termite species with different diets. Termite species are arranged by the degree of plant degradation in the diet. **(B)** Principal component analysis of community similarities of termites with different diet. Mn: *Macrotermes natalensis*, Od: *Odontotermes* sp., Nc: *Nasutitermes corniger*, Aw: *Amitermes wheeleri*, Mp: *Microcerotermesparvus*, Co: *Cornitermes* sp., Th: *Termes hospes*, Nt: *Neocapritermes taracua*, Cu: *Cubitermes ugandensis*.

### Fungus- and plant cell wall-targeting enzymes

To compare the functional capacity for fungal and plant cell wall degradation, we identified genes belonging to GH families in the CAZy database (Cantarel et al. 2009; Lombard et al. 2014), and classified the genes by their substrate target and thus putative enzyme function by assigning EC numbers (Table 2, S3). Principle Component Analysis (PCA) of GH compositions (Figure 2B) support that gut microbial enzyme capacities are similar between termites with similar diets. This link was also apparent when evaluating relative abundances of GH families coded for by the gut bacteria (Figure 2C). Fungus-growing termite guts were thus systematically different in GH family composition to the guts of termites feeding on plant biomass (Figure 2B, C), reflecting the radical differences in cell wall composition of plant and fungus material.

**Figure 2.**
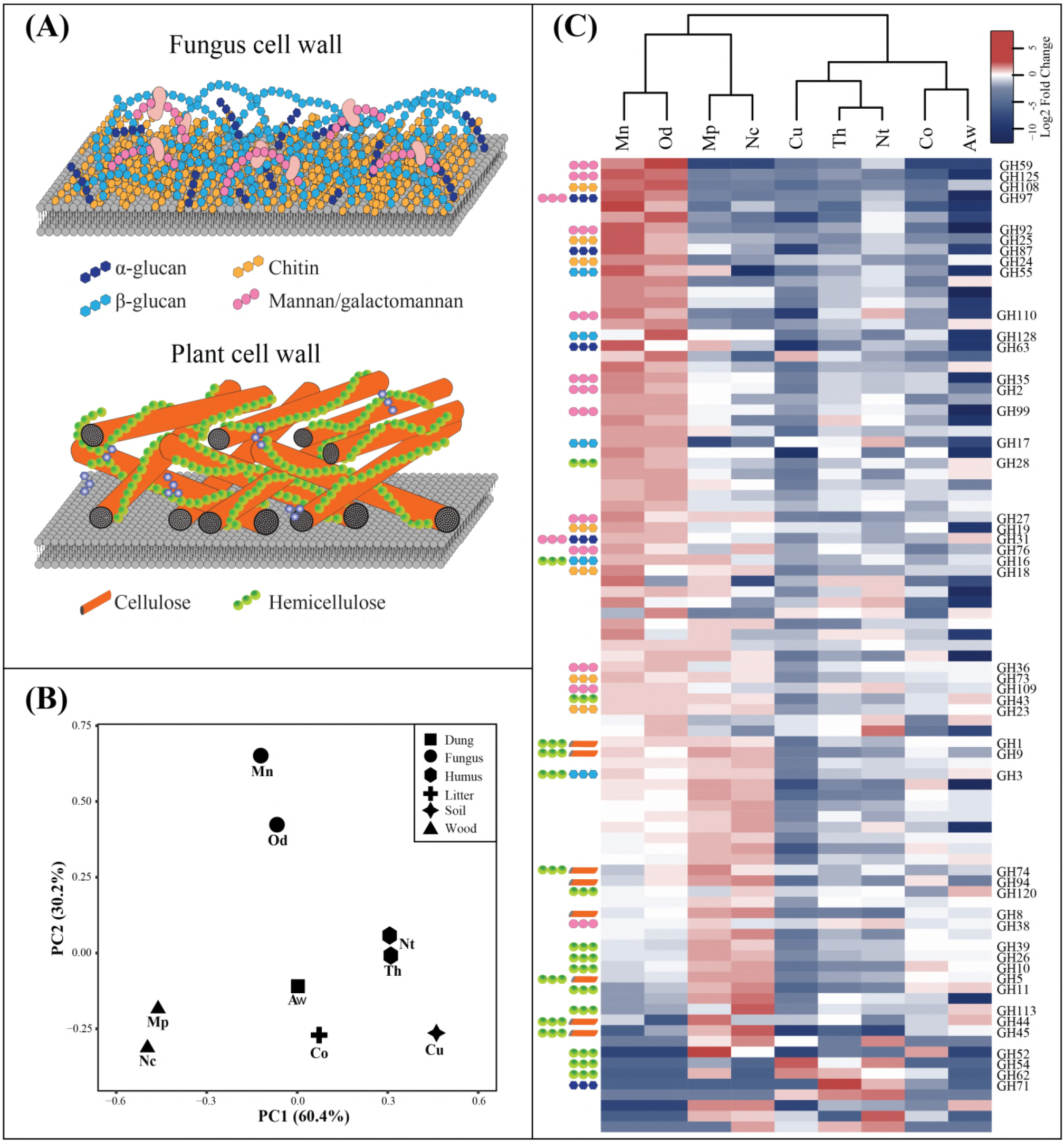
**(A)** Simplified schematic of fungal (top) and plant (bottom) cell wall structures and targets of enzymes identified in the metagenomes. **(B)** Principal component analysis (PCA) of relative abundances of carbohydrate-active enzymes in worker gut metagenomes; shapes represent termite feeding groups. Mn: *Macrotermes natalensis*, Od: *Odontotermes* sp., Nc: *Nasutitermes corniger*, Aw: *Amitermes wheeleri*, Mp: *Microcerotermes parvus*, Co: *Cornitermes sp.*, Th: *Termes hospes*, Nt: *Neocapritermes taracua*, Cu: *Cubitermes ugandensis.* **(C)** Relative abundances of enriched (red) and contracted (blue) GH families across the nine metagenomes shown as Log2 fold changes. The symbols on the left of the heatmap follow the legend from Figure 2A, and indicate which predicted substrates that are targeted by the enzymes by members of a given GH family, and family names of GH families with the highest average abundances across all metagenomes are indicated on the right.

GH enzymes targeting fungal cell wall components were enriched in fungus-growing termites, with the most marked differences being for GH125 (34-fold) and GH92 (14-fold) that likely target α-1,6-and 1,2-mannoside (Figure 2C; Table S2). Several β-1,3-glucan-targeting families (GH17, GH128, GH55, and GH87), the chitin-targeting families GH18 and GH19, and lysozyme families with chitin-degradation activities (GH23, GH73, GH24, and GH25), showed 2- to 12-fold enrichment in fungus-growing termites (Figure 2C; Table S2). Similarly, enzymes target galactans that likely belong to fungal cell wall were enriched: GH families targeting β-galactosidase (GH59, GH35, and GH2), along with α-galactosidase and α-N-acetylgalactosaminidase (GH110, GH27, and GH97), showed 3- to 36-fold enrichment compared to non-fungus growing termites. In contrast, and as expected, many GH families encoding genes targeting plant cell wall components were low in relative abundance in fungus-growing termites but higher in wood feeders (Figure 2C; Table S2). Cellulose- (GH94, GH10, GH5) and hemicellulose-targeting families (GH74, GH120, GH39, GH26, GH10, GH11 etc.) were 3- to 5-fold more abundant in wood feeders than in other termites. Interestingly, laccases (EC 1.10.3.2) were only detected in wood feeding termites (Table S3). Lytic polysaccharide monooxygenase (LMPO) were only detected in *Cu. ugandensis*, where they, however, were present only in very low abundance (Table S3).

### The bacteria encoding the most abundant glycoside hydrolase families

To gain further insight into the functional contribution of gut microbiota members to fungal digestion, we grouped enzymes in GH families by fungal cell wall components targeted and bacteria of origin (Table 2; Figure 3). The bacterial orders Clostridiales and Bacteroidales contributed most fungal cell wall-degrading enzymes in fungus-growing termite guts (78%), while Spirochaetales contributed most of these enzymes in the wood-feeding termites (62%; Figure 3). Lysozymes targeting chitin was only abundant in fungus-growing termite gut metagenomes (Table 2; Figure 3) and mainly, coded for by members of the Clostridiales. α-mannanase, β-glucanase, and galactosidase targeting mannan/galactomannan and β-glucan in the fungal cell wall, respectively, were also abundant in fungus-growing termite gut, with the majority of these enzymes (60%) being coded for by members of the Bacteroidales (Figure 3), while chitinases and β-N-acetylhexosaminidases were produced by both Clostridiales and Bacteroidales and, to a lesser extent, Spirochaetales in wood feeders. In the remaining termite species, most of the fungal cell wall-degrading enzymes were contributed by Clostridiales, Bacteroidales, and Spirochaetales (17%-92%), but the overall abundance of these enzymes was far lower than in fungus-growing termites and wood feeders (Table 2; Figure 3).

**Figure 3.**
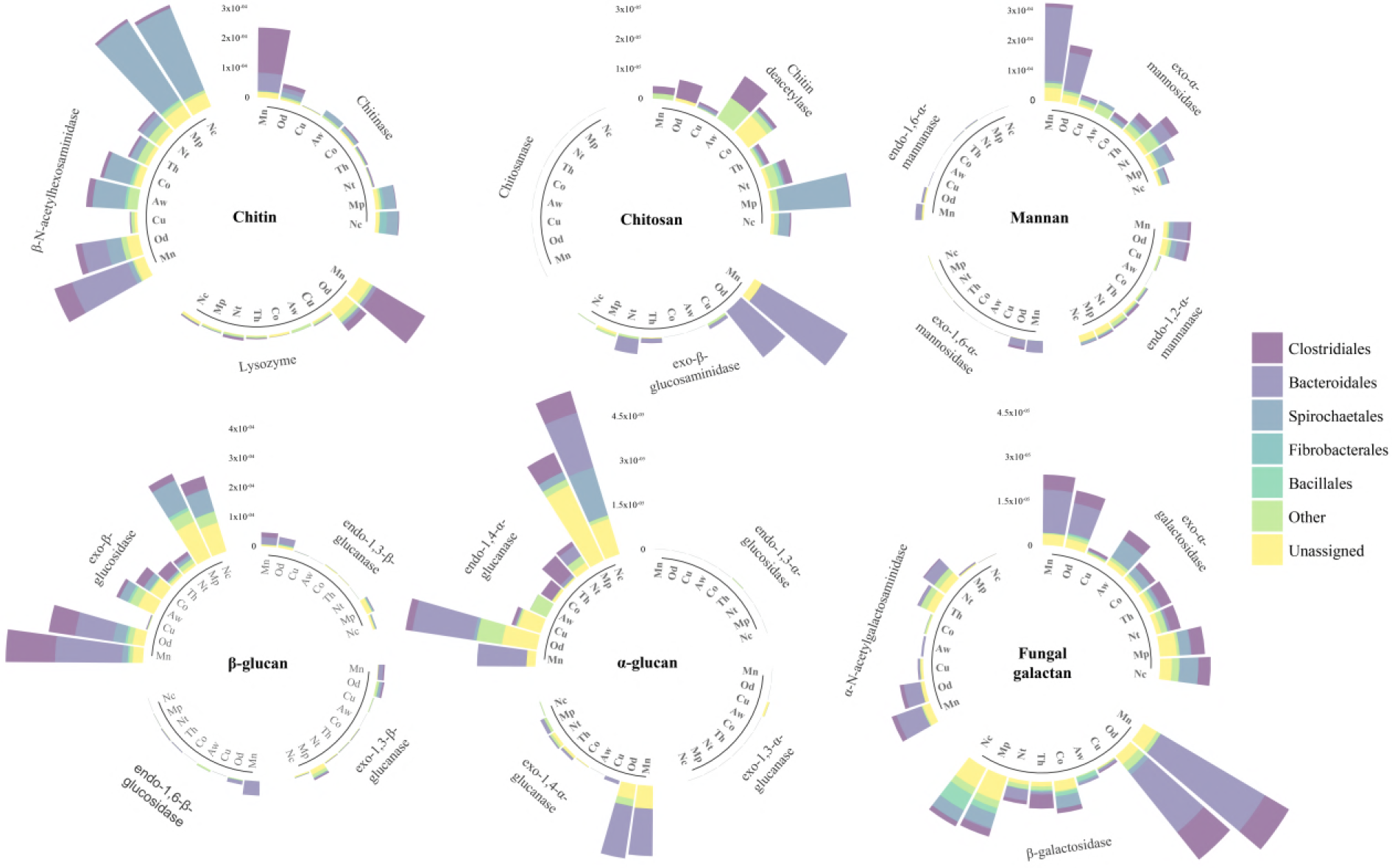
The relative abundance of fungal cell wall degrading enzymes encoded by the top five most abundant bacterial orders in the nine termite species. Mn: *Macrotermes natalensis*, Od: *Odontotermes* sp., Nc: *Nasutitermes corniger*, Aw: *Amitermes wheeleri*, Mp: *Microcerotermes parvus*, Co: *Cornitermes* sp., Th: *Termes hospes*, Nt: *Neocapritermes taracua*, Cu: *Cubitermes ugandensis*.

### Distinct protease profiles in termite gut metagenomes

We identified proteases in termite gut metagenomes and classified them by catalytic types to investigate the proteolytic potential of gut communities. Proteases were most abundant in fungus-growing termites, which exhibited a 1.4-4 fold enrichment in all catalytic types (Figure 4A), followed by *Mi. parvus*, *Na. corniger* (wood-feeders) and *A. wheeleri* (dung-feeder) (Figure 4B; Table S4). One of the most distinct differences we observed was for glutamic peptidases and mixed peptidases, which were enriched only in the guts of fungus-growing termites. The wood feeder *Cu. ugandensis* displayed the lowest protease abundance, followed by the litter-feeder *Cornitermes* sp. As observed for fungal and plant cell wall-degrading enzymes, the most abundant taxa were also predicted to encode for most of the proteases (Figure 4A). Clostridiales and Bacteroidales, Clostridiales and Spirochaetales, and Spirochaetales were the main contributors encoding proteases within the metagenomes of fungus-feeding, dung-feeding, and wood-feeding termites, respectively.

**Figure 4.**
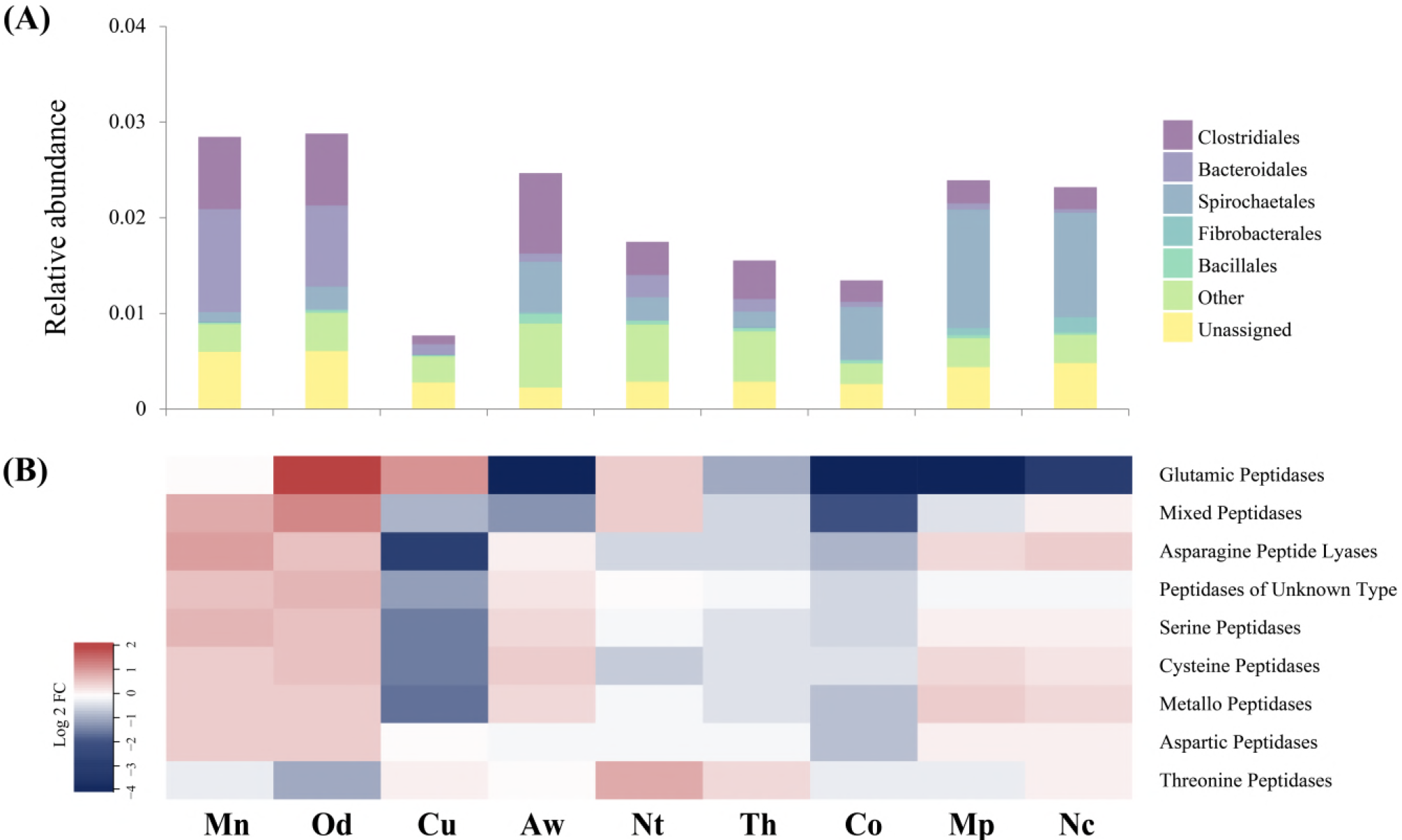
**(A)** The relative abundances of proteases encoded for by the five most contributing bacterial orders across the nine termite species. **(B)** A heatmap of the relative abundances of the most abundant proteases across the nine species shown as Log2 fold change (red: enriched and blue: contracted). Mn: *Macrotermes natalensis*, Od: *Odontotermes* sp., Nc: *Nasutitermes corniger*, Aw: *Amitermes wheeleri*, Mp: *Microcerotermes parvus*, Co: *Cornitermes* sp., Th: *Termes hospes*, Nt: *Neocapritermes taracua*, Cu: *Cubitermes ugandensis*.

## Discussion

Diet is a major driver of taxonomic and functional composition of gut microbial communities (David et al. 2014 Perez-Cobas et al. 2015; Bodawatta et al. 2018). Our characterisation of cell wall degrading enzyme and protease enzymes from higher termites confirm that the functional profiles of gut microbial communities are tight linked with termite diet, consistent with previous work that has focused on community compositions (Otani et al. 2014, 2016; Mikaelyan et al. 2015), and preliminary functional implications of the shift in microbiota structure after the origin of fungus farming in the termite sub-family (Liu et al. 2013; Poulsen et al. 2014). The gut microbiota thus not only functionally complements final plant biomass decomposition by providing oligosaccharide-targeting enzymes (Liu et al. 2013; Poulsen et al. 2014), but also provides key enzymes for the digestion of fungal biomass. The mycolytic potential of the gut microbiota of the South African *Odontotermes* sp. and *Ma. natalensis* (this study) was comparable *O. yunnanensis* from Southwest China (Liu et al. 2013) on, suggesting conserved functions across space and time, and highlighting the robust link between taxonomic and functional ties of the intimate interactions between gut community members and termite hosts.

Fungus-growing termite gut mycolytic enzymes were primarily coded for by Clostridiales and Bacteroidales, which dominate the core gut microbiota of the Macrotermitinae (Hongoh et al. 2005, Hongoh 2010; Dietrich et al. 2014; Otani et al. 2014; Mikaelyan et al. 2015). The ancestor of the Macrotermitinae likely had a lower termite bacterial gut microbiota (but without protists) (Brune 2014; Brune and Dietrich 2015). Fungiculture exposed the gut microbiota to higher amounts of fungal biomass including proteins than to the ancestral lignocellulolytic diet, resulting in a gut microbiota that converged to become more similar to those observed in extant cockroaches (Dietrich, Kohler, and Brune 2014; Otani et al. 2014). This is likely a product of several factors. First, bacterial strains present in the Macrotermitinae ancestor that could utilise a fungal diet were likely selected for and consequently increased in relative abundance (e.g., *Alistipes*, *Dysgonomonas*, and members of the *Ruminococcaceae*) (Dietrich et al. 2014; Otani et al. 2014). Many mycolytic microbes were conceivably already present in termite guts prior to the evolution of fungiculture, since termites feed on partially degraded plant substrates containing fungal biomass. Second, bacteria contributing to the breakdown of lignocellulose in the Macrotermitinae ancestor were likely outcompeted/selected against (e.g., the genus *Treponema)*. Third, novel lineages adopted from other termites or the environment were likely co-opted when fungiculture evolved. This is consistent with recent work demonstrating rampant horizontal transmission of gut bacterial lineages associated with termites (Bourgoignon et al. 2018). Collectively, this led to a gut microbiota enriched in mycolytic (e.g., α-mannanases, β-glucanases, chitinases and proteases) and reduced in lignocellulolytic (e.g., cellulase, cellobiohydrolase, hemicellulose and laccase) enzymes.

In sharp contrast to the fungus growers, lignocellulose-degrading enzymes dominated the gut microbiota of wood-feeding termites (*Na. corniger* and *Mi. parvus*), as expected from the requirement for the breakdown of recalcitrant plant components, which in fungus farmers are handled by *Termitomyces* (Poulsen et al. 2014; although lignin cleavage may be initiated during the first gut passage: Li et al. 2017). Interestingly, however, the relatively high abundance of chitinolytic enzymes of Spirochaetales origin (mostly species of the genus *Treponema*) in wood-feeding termites suggests that the decaying wood these species feed on, harbour fungal biomass that gut bacteria or the termite host may utilise. However, it is also conceivable that fungal cell wall-degrading enzymes, such as β-1,3-glucanase, serve to cleave fungal cell wall to protect against fungal infections (cf., Rosengaus et al. 2014). Experimental work targeting their expression in the presence of fungal pathogens, ideally combined with varying fungal biomass diet content, have the potential to shed light on their relative defensive and dietary roles.

Proteases were most abundant in fungus-growing termites and mostly coded for by the bacterial taxa contributing mycolytic enzymes. The presence and high abundance of nearly all types of proteases reflects a general and broad proteolytic capacity of the fungus-growing termite gut microbiota that is inevitably related to the protein-rich fungal diet. The presence of proteases in dung and wood feeders could be explained by the presence of fungal biomass in the substrate ingested by these termites. This is consistent with the low abundance of these enzymes in termites feeding on soil and humus with much lower protein content. The only under-represented proteases in fungus-growing termites was threonine peptidases, which has comparable activities as serine and cysteine peptidases (Powers et al. 2002). The absence of these enzymes could thus be compensated for by the presence of other protease families; however, more work will be needed to test what these enzymes indeed target.

Similarities in host diet have been shown to drive convergence in the functional potential of gut microbes in other organisms (Muegge et al. 2011; Delsuc et al. 2014). Selection for particular physiological traits may, however, not necessarily be directly linked to specific phylogenetic groups of microbes. Exploring communities associated with diverse fungus-growing hosts (and their associated fungi) would allow us to explicitly test for convergent evolution of chitinolytic microbial communities, even if these were likely comprised by different microbial consortia. A number of other insects utilise fungus material as a nutrient source, including fungus-growing ants (Schultz and Brady 2008; Schiøtt et al. 2010), some *Drosophila* species (Jaenike and James 1991), the Malaysian mushroom-harvesting ant *Euprenolepis* (Witte and Maschwitz, 2008), *Sirex* wood wasps (Madden and Coutts, 1979; Hajek et al. 2013), and ambrosia beetles (Batra 1966; Hulcr and Cognato 2010; De Fine Licht and Biedermann 2012). Different bacteria are likely to be involved, but predictions would be that predominantly fungal diets should select for microbial communities with comparable mycolytic capacities. Recent work on mycophagous *Drosophila* supports that this may be so. Bost et al. (2018) demonstrated that gut bacteria in mycophagous *Drosophila* are implicated in fungal cell wall metabolism, with cysteine and methionine metabolism enzymes originating from Bacteroidetes, Firmicutes, and Proteobacteria gut microbiota members; phyla that are also abundant and mycolytic in fungus-growing termites. The convergent prevalence of specific protease-producing Firmicutes and Bacteroidetes taxa suggests that they were selected for high-protein host diets, consistent with findings in humans (Eckburg et al. 2005) and pigs (Leser et al. 2002), and likely contributing to the observed convergence in fungus-growing termite and cockroach gut metagenomes (Dietrich et al. 2014; Otani et al. 2014; Schauer et al. 2012).

The prevalence of microbial communities with ample mycolytic capacities in the guts of fungus-growing termite species supports that the shift to a fungal diet was associated with both a functional and compositional shift in gut microbial communities at the onset of fungiculture in termites. The enrichment of GH families encoding fungal cell wall-degrading enzymes and proteases indicates adaptations to the decomposition a fungal diet, consistent with this capacity being absent or less in termites with predominantly plant-based diets. An exception is wood-feeding termites, for which wood-degrading fungi also may comprise an appreciable component of the termite diet. Further work will be needed to elucidate whether the functional capacities of the gut microbiota reflect the amount of fungal biomass in the diet and potential differences in dietary properties of the fungal species fed on. Furthermore, taxonomy assignments were reference-dependent and limited by the closest match present in the databases, with 10-60% of the enzymes identified per metagenome remaining unclassified and most of the classified enzymes remaining unidentified at the genus level, so to improve our understanding of digestive function of gut microbes, further classification of the taxonomic and functional properties of digestive enzymes and nutrient composition of termite diets are needed.

## Material and Methods

### Odontotermes *sp. collection*

Termites from an *Odontotermes* sp. colony (code: Od127) were collected at the Experimental Farm of the University of Pretoria, South Africa (-25.742700, 28.256517). The species identity of this colony had been previously established as by mitochondrial gene COII barcoding (Otani et al. 2014). Fifty old major workers were sampled, and entire guts were dissected and pooled in a 1.5ml Eppendorf tube and stored at -80°C until DNA extraction.

### Gut microbiota DNA extraction

Guts were ground in liquid nitrogen, after which DNA was extracted using the Qiagen Animal Tissue Mini-Kit (Qiagen, Hilden, Germany) according to the manufacturer’s description, with the exception that a chloroform extraction step was followed by incubation with protease K. After proteinase K digestion, one volume chloroform/isoamyl alcohol (24/1) was added; tubes were incubated for 15min on a slowly rotating wheel, and centrifuged at 3,000 g for 10 min. The supernatant was transferred to spin columns and the remainder of the manufacturer’s protocol was followed. The quality and purity of samples were determined using NanoDrop^®^ (Thermo Scientific, Wilmington, USA).

### Metagenome sequencing and assembly

DNA was sheared to ~350bp fragments, end-repaired, A-tailed, and ligated with Illumina paired-end adaptors (Illumina). The ligated fragments were selected from the desired size on agarose gels and amplified by LM-PCR, and libraries were sequenced with 150bp read lengths on an Illumina HiSeq2500. The quality of raw sequencing reads was assessed before assembly. Reads containing the adaptor, more than 10% N or more than 50% low quality bases (Q-score≥ 5), were removed. To exclude sequences from *Termitomyces* and the termite hosts, quality-controlled reads were mapped to the *Ma. natalensis* and *Termitomyces* genomes (Poulsen et al. 2014) with Burrows-Wheeler Aligner v0.7.15 (Li and Durbin 2010) BWA-MEM algorithm; any aligned reads were filtered.

Clean reads were assembled by IDBA-UD v1.1.2 (Peng et al. 2010; 2011) with an iterative set up from k-mer size of 19 to 99 at step of 10 (--pre_correction --mink 19 --maxk 99 --step 10). Unassembled reads were picked out by mapping reads back to the initial assembly and assembled separately with the same set up. Redundancies of sequences from the same organism within the metagenome were removed by clustering all contigs at 95% identity with CD-hit v4.6.6 (Li and Godzik 2006) and only the longest contig per cluster was kept. For comparison, high-quality reads of *Ma. natalensis* old major worker gut microbiota from Poulsen et al. (2014) were also reassembled using the same procedure. Genes in both assemblies were predicted by Prodigal v2.6.3 (Hyatt et al. 2010) with metagenomics parameters (-c–p meta).

### Non-fungus growing termite gut metagenomes

We obtained seven published non-fungus growing termite gut metagenomes. These were a dung feeder: *Amitermes wheeleri* (He et al. 2013); two wood feeders: *Nasutitermes corniger* and *Microcerotermesparvus;* a litter feeder *Cornitermes* sp.; two humus feeders: *Termes hospes* and *Neocapritermes taracua;* and the soil feeder *Cubitermes ugandensis* (Rossmassler et al. 2015). Contigs and protein coding genes were downloaded from JGI IMG/M (Table 1).

**Table 1.** Summary of metagenome data.

| Termite species | Diet | Reference | Sample data size (Gbp) | Accession No. | Assembly size (Mbp) | Assembly N50 (bp) | Number of genes |
| --- | --- | --- | --- | --- | --- | --- | --- |
| *Odontotermes* sp. | Fungus | NA | 8.6 | PRJNA476694 | 797.15 | 1,133 | 753,265 |
| *Macrotermes natalensis* | Fungus | Poulsen et al. 2014 | 8.5 | PRJNA193472 | 498.71 | 1,569 | 486,207 |
| *Amitermes wheeleri* | Dung | He et al. 2013 | 0.3 | IMG:2030936000 | 139.3 | 512 | 321,461 |
| *Nasutitermes corniger* | Wood | Rossmassler et al. 2015 | 46.8 | IMG:3300002119 | 634.52 | 980 | 1,338,688 |
| *Microcerotermes parvus* | Wood | Rossmassler et al. 2015 | 43.2 | IMG:3300002449 | 712.35 | 945 | 1,419,719 |
| *Cornitermes* sp. | Litter | Rossmassler et al. 2015 | 45.9 | IMG:3300002450 | 131.61 | 519 | 2,945,692 |
| *Termes hospes* | Humus | Rossmassler et al. 2015 | 34.2 | IMG:3300002469 | 1212.37 | 424 | 2,975,319 |
| *Neocapritermes taracua* | Humus | Rossmassler et al. 2015 | 28.3 | IMG:3300002505 | 885.58 | 454 | 2,115,406 |
| *Cubitermes ugandensis* | Soil | Rossmassler et al. 2015 | 32.0 | IMG:3300002127 | 1121.61 | 475 | 3,061,751 |

**Table 2.**
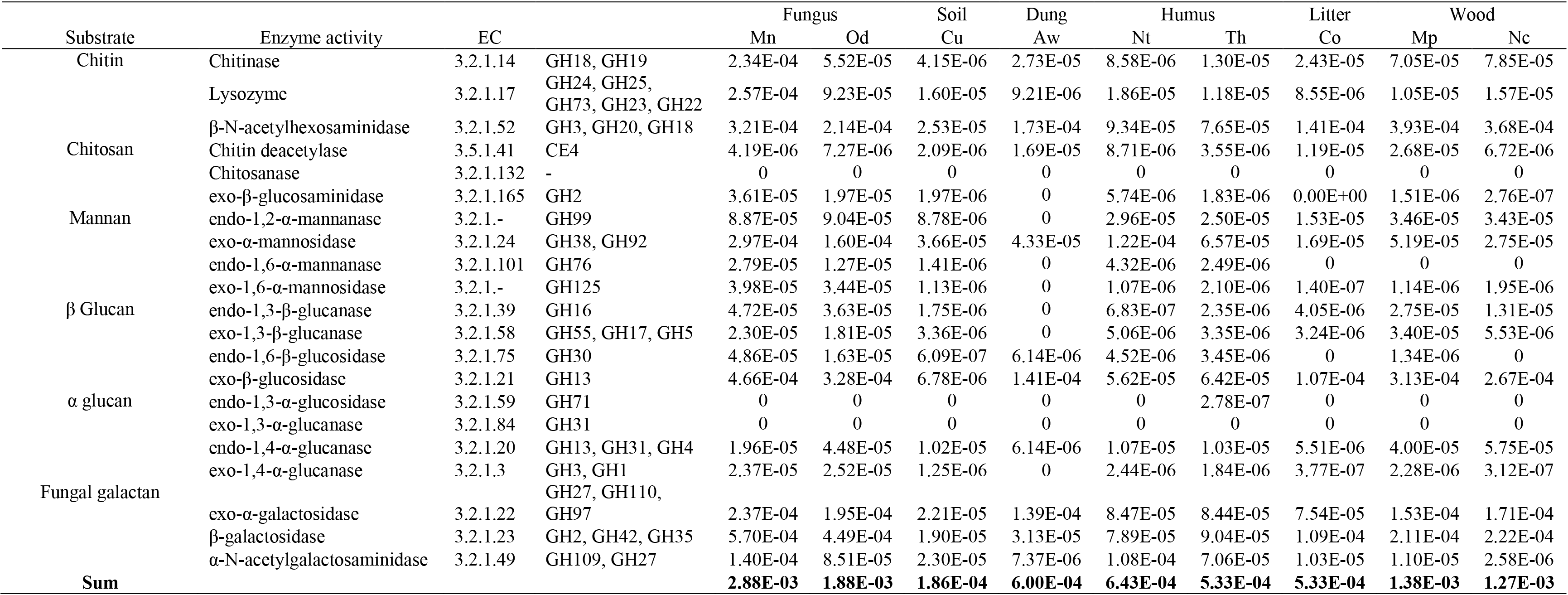
Relative abundance of fungal cell wall degrading enzymes in the gut metagenomes of nine termites with different diets. Mn: *Macrotermes natalensis*, Od: *Odontotermes* sp., Nc: *Nasutitermes corniger*, Aw: *Amitermes wheeleri*, Mp: *Microcerotermes parvus*, Co: *Cornitermes* sp., Th: *Termes hospes*, Nt: *Neocapritermes taracua*, Cu: *Cubitermes ugandensis.* For the full results, see Table S2.

### Relative abundances of contigs within metagenomes

Clean reads of fungus-growing termites gut microbiota were mapped to the contigs by Bowtie v2.2.9 (Langmead and Salzberg 2012), with subsequent masked duplication taking the best match for each read. Contig coverage was estimated by the length and the number of mapped reads per contigs, after which the relative abundance of each contig was calculated as the coverage of the contig divided by the sum of coverage of all contigs (Qin et al. 2010). For non-fungus growing termites, coverage information of contigs was obtained from JGI (Nordberg et al. 2014; Grigoriev et al. 2012) and the relative abundances were estimated in as described for fungus-growing termites.

### Metagenome taxonomic assignment

Taxonomic assignment of protein-coding genes was carried out using Diamond (Buchfink et al. 2015) alignment against the NR database in NCBI. Alignments with e-values >1e-5 and sequences with identities <30% were removed. Taxonomic information of the top hit was assigned to each gene. Contigs referring to taxonomical levels were determined by a modified lowest common ancestor (LCA)-based algorithm implemented in MEGAN (Huson et al. 2007). Taxonomic classification supported by less than 10% of the genes on each contig were first filtered, and the lowest common ancestor (LCA) for the taxonomic classification of the rest of the genes were assigned to the contig. The relative abundance of contigs belonging to the same taxonomic group was summed up to represent the taxonomic abundance of that taxonomic group in the microbiota. Differences in taxonomic abundances at the phylum level for metagenomes were compared and visualized by Principle Component Analysis (PCA) using R v3.3.2 (R Core Team 2013).

### Carbohydrate-active enzyme analysis

We identified genes encoding CAZymes by searching against CAZyme hidden Markov profiles in the dbCAN database (Yin et al. 2012) using HMMer v3.1b (e ≤ 1e-5) (Eddy et al. 1994) and peptide pattern recognition using HotPep (Busk et al. 2013; 2014; 2017). CAZyme family classification supported by both methods was assigned to genes. Genes for which the two methods gave inconsistent results were subjected to an additional Pfam protein domain search using HMMer (e ≤ 1e-6) to determine the CAZyme family. CAZyme compositional differences between termite species were evaluated using PCA in R. Enzyme functions of genes in each CAZyme family were determined by BLASTp searches (e ≤ 1e-5) against the ExPASy enzyme records, CCD searches against COG database (Marchler-Bauer et al. 2017) and GhostKOALA searches using the KEGG database online tool (Kanehisa et al. 2016). Genes coding for enzymes related to fungal and plant cell wall degradation were selected and their substrate targets and bacterial taxonomy assigned and manually checked.

### Protease identification and analyses

Protein sequences of annotated genes in metagenomes were aligned to MEROPS database (Rawlings et al. 2018) by BLASTp (e ≤ 1e-5) to identify proteases. Genes that matched at least 50% of the sequences in the database were selected. The catalytic type and protease family classification of genes were annotated based on their most similar matches in the MEROPS database, and bacterial taxonomy was assigned and manually checked as described for the CAZyme analysis.

## Data deposition

Clean reads and metagenome assembly have been submitted to the SRA and GenBank under BioProject accession numbers PRJNA476694 and PRJNA193472.

## Acknowledgements

We thank Z. Wilhelm de Beer, Michael J. Wingfield, and the staff and students at the Forestry and Agricultural Biotechnology Institute, University of Pretoria, for hosting field work and contributed to the research environment, and Christine Beemelmanns, René Benndorf, Saria Otani, Margo Wisselink, Sabine M. E. Vreeburg, Duur K. Aanen, Lennart van de Peppel, Nina Kreuzenbeck, and Victoria L. Challinor for help with excavations. This study was funded by the CAPES Foundation, Ministry of Education of Brazil (grant BEX: 13240/13-7) to RRdC and a Villum Kann Rasmussen Young Investigator Fellowship (VKR10101) to MP.

## Author Contributions

HH and RRdC designed the experiments and analyses, collected material in the field, extracted DNA, sent samples for sequencing, and drafted the first version of the figures, tables, and manuscript. HH carried out the assembly of the metagenome and the functional analyses. BP and LL carried out HotPep analyses and contributed to data interpretation. MS contributed to data analyses and interpretations. MP supervised HH and RRdC, helped design the study, contributed with comments on analyses and the first versions of figures, tables, and text. All authors contributed to writing the manuscript.

## Supplementary Material

**Table S1.**
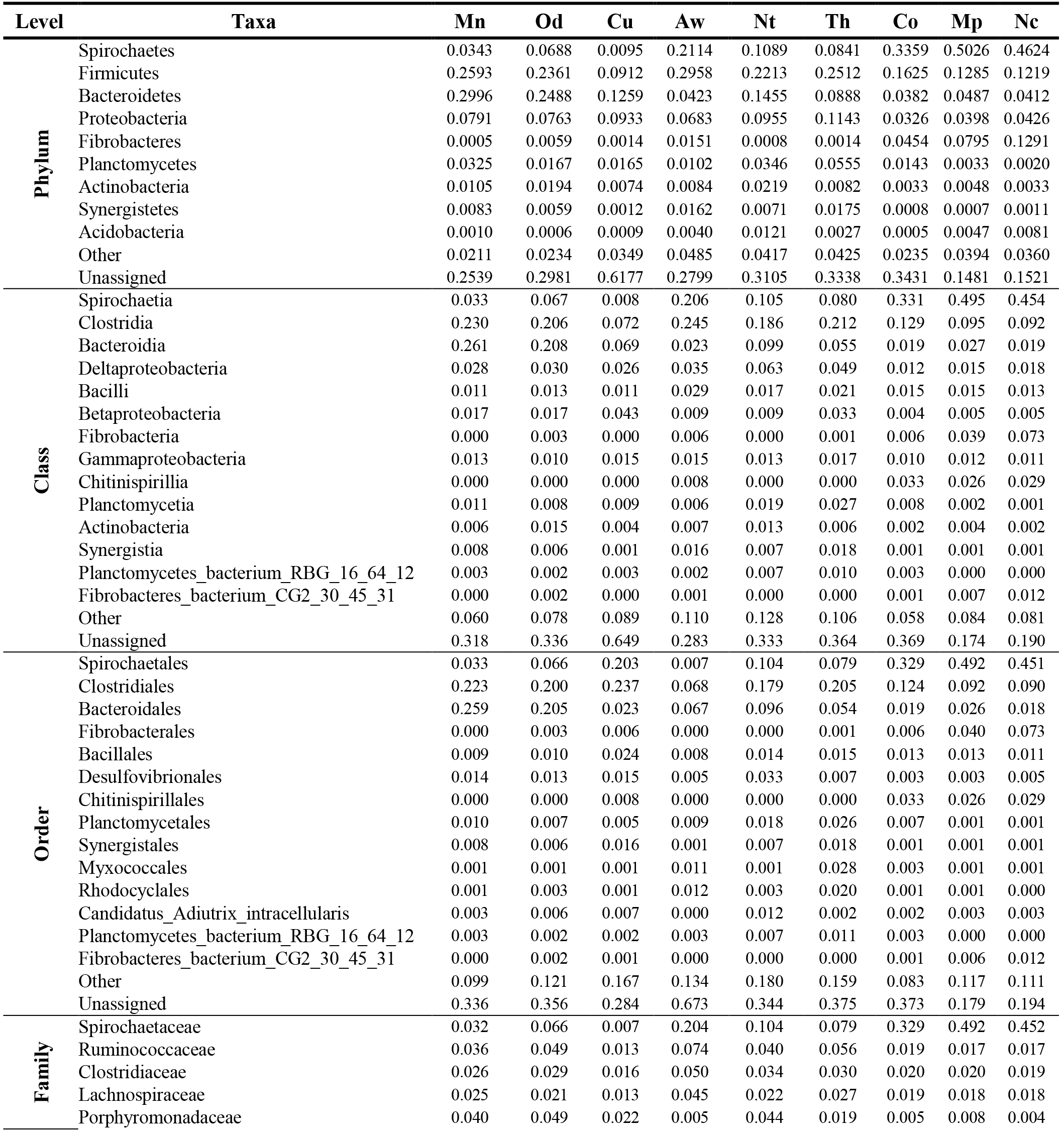
Bacterial community composition of metagenome contigs of gut metagenomes of nine termites with different diets at the phylum, class, order, family, and genus levels. Taxonomy classifications with more than 0.1% of the total abundance are shown. Mn: *Macrotermes natalensis*, Od: *Odontotermes* sp., Nc: *Nasutitermes corniger*, Aw: *Amitermes wheeleri*, Mp: *Microcerotermes parvus*, Co: *Cornitermes* sp., Th: *Termes hospes*, Nt: *Neocapritermes taracua*, Cu: *Cubitermes ugandensis*.

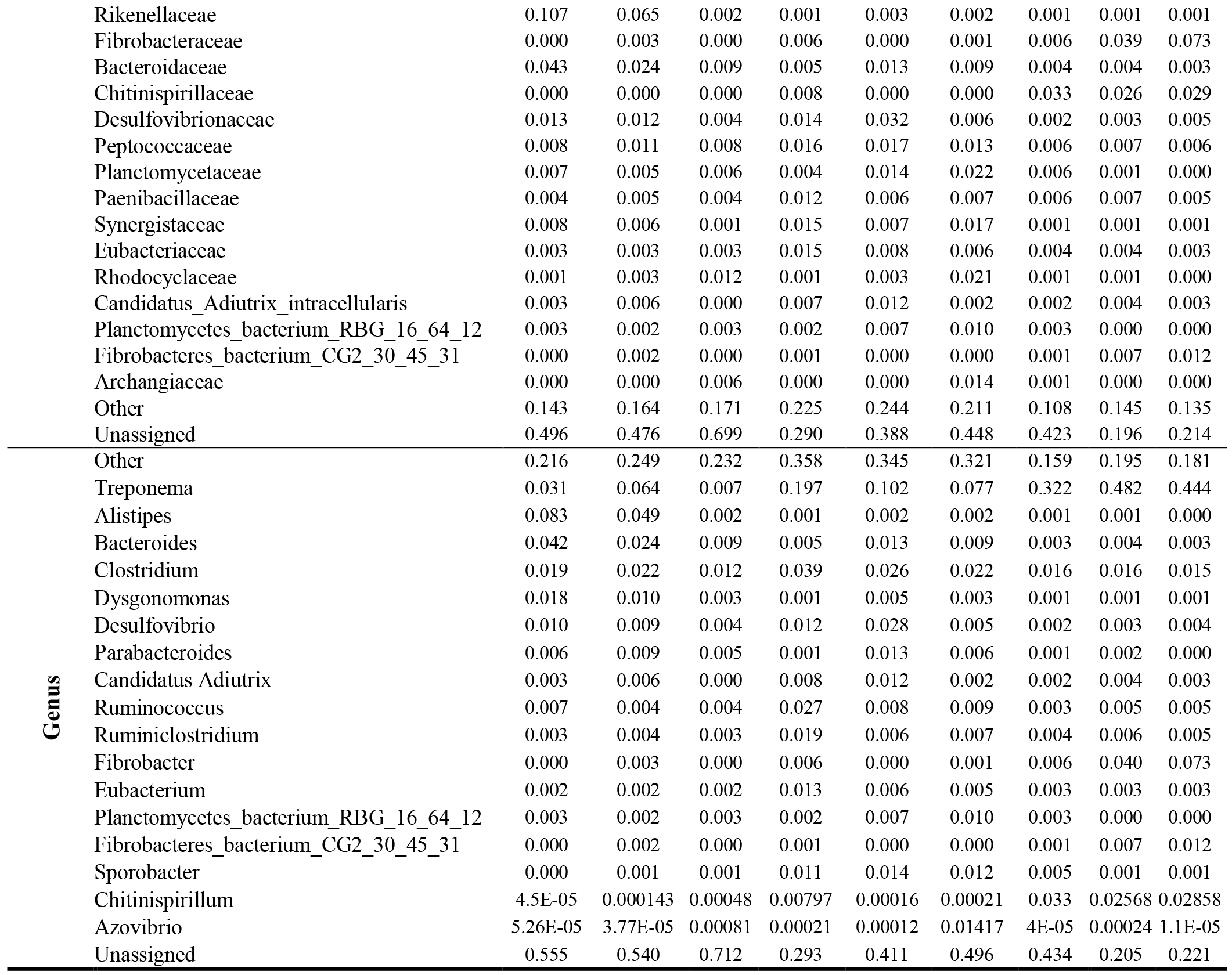

**Table S2.**
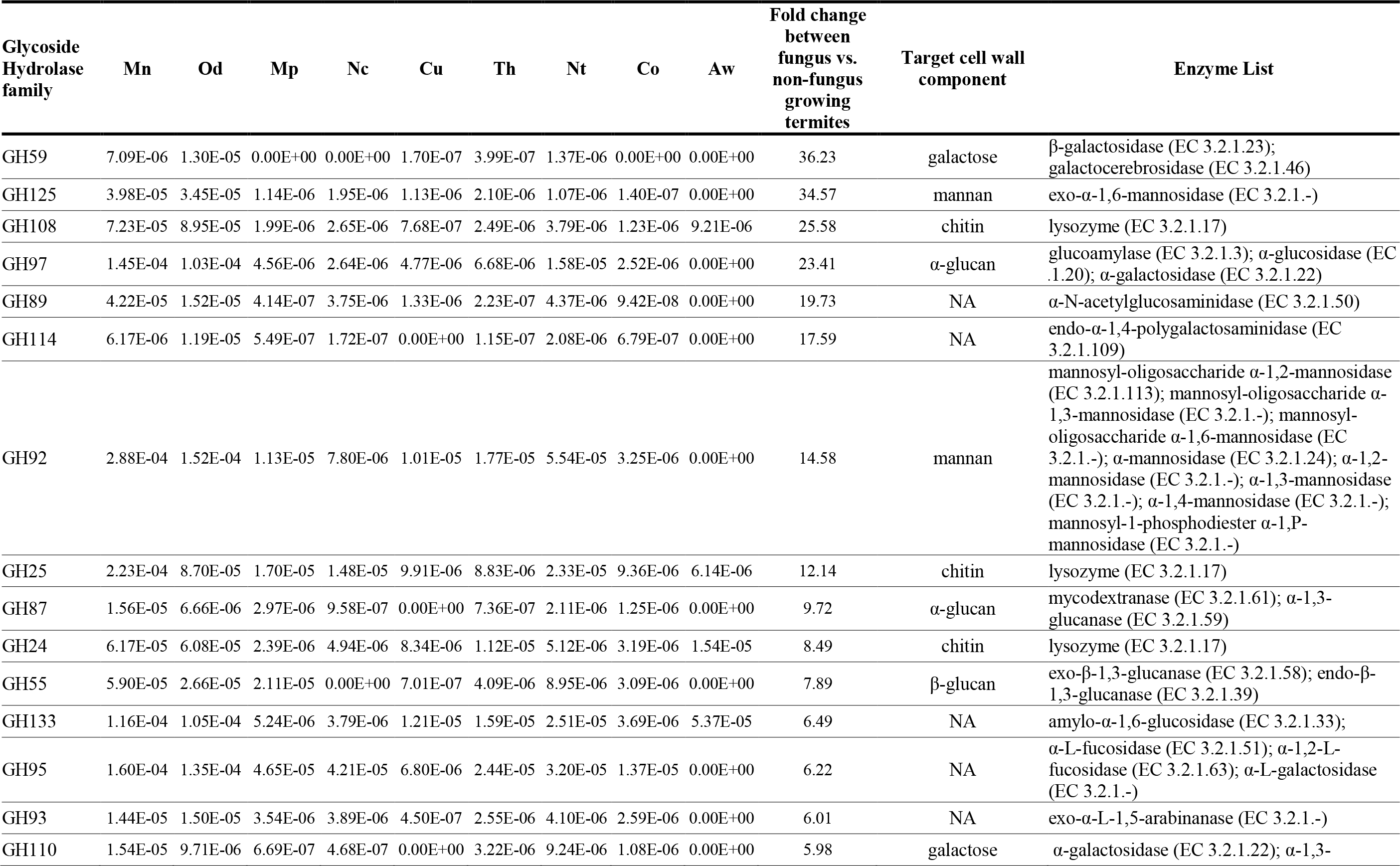
Relative abundance of glycoside hydrolase families in gut metagenomes of nine species of termites with different diets. Mn: *Macrotermes natalensis*, Od: *Odontotermes* sp., Nc: *Nasutitermes corniger*, Aw: *Amitermes wheeleri*, Mp: *Microcerotermes parvus*, Co: *Cornitermes* sp., Th: *Termes hospes*, Nt: *Neocapritermes taracua*, Cu: *Cubitermes ugandensis*.

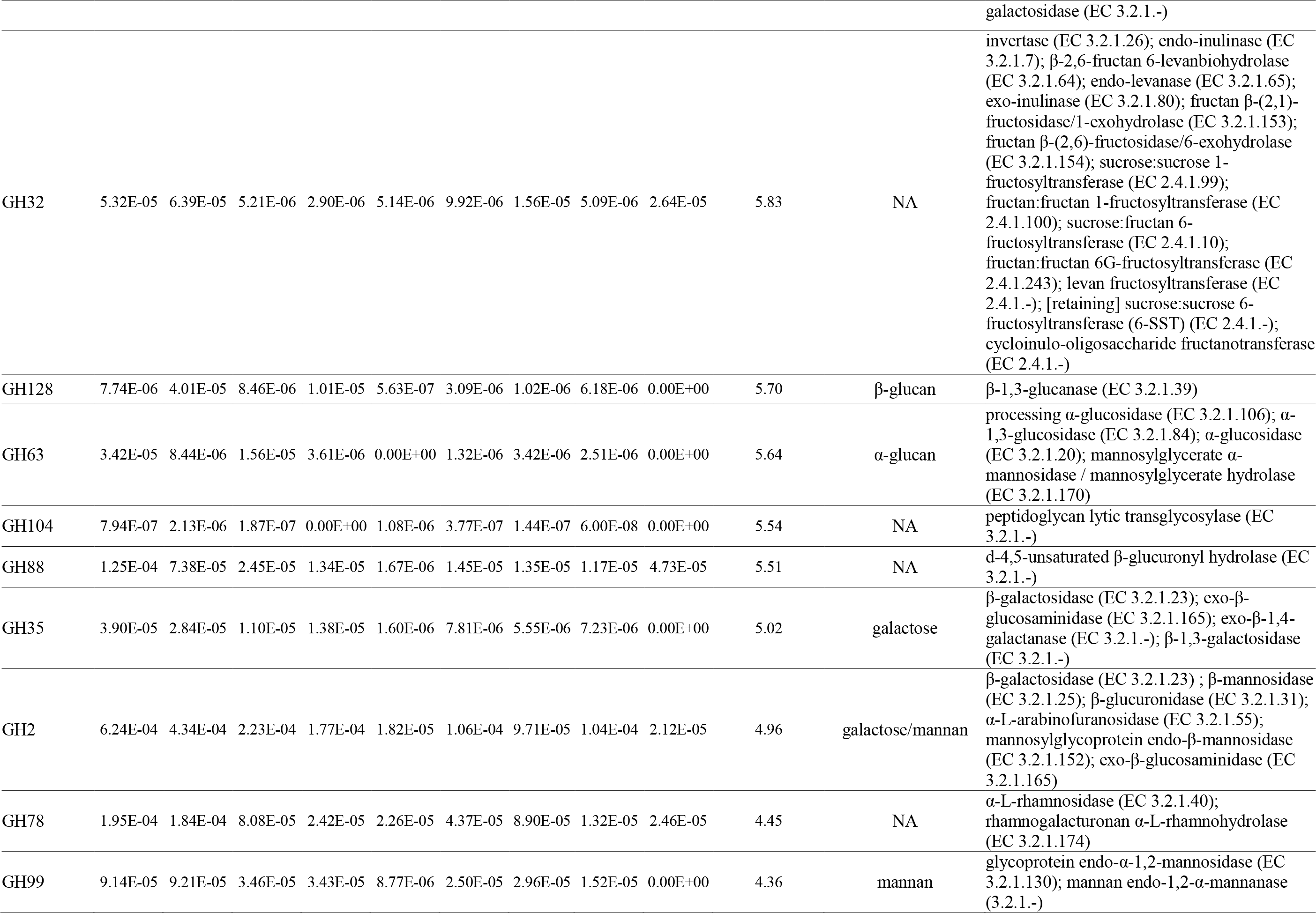

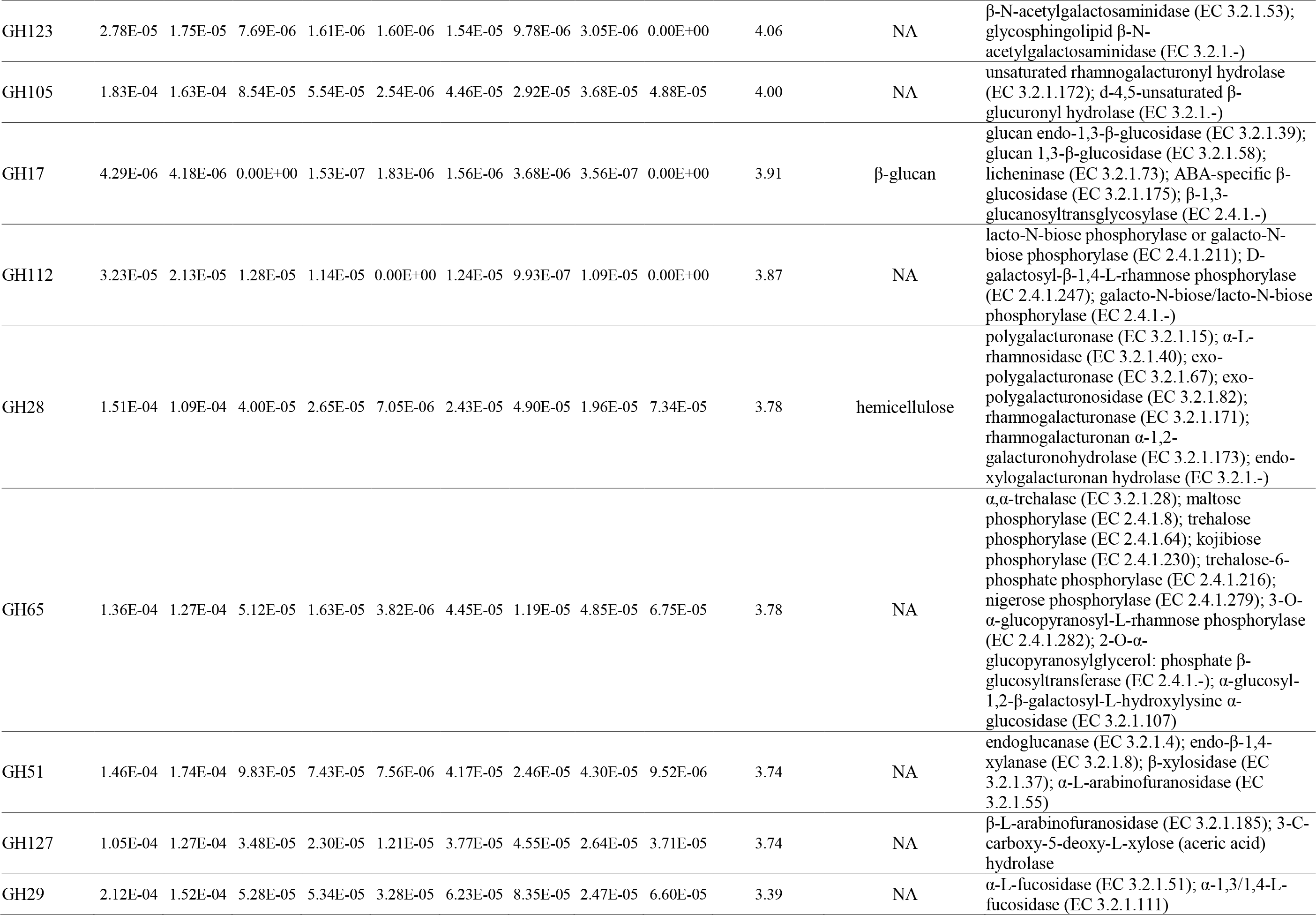

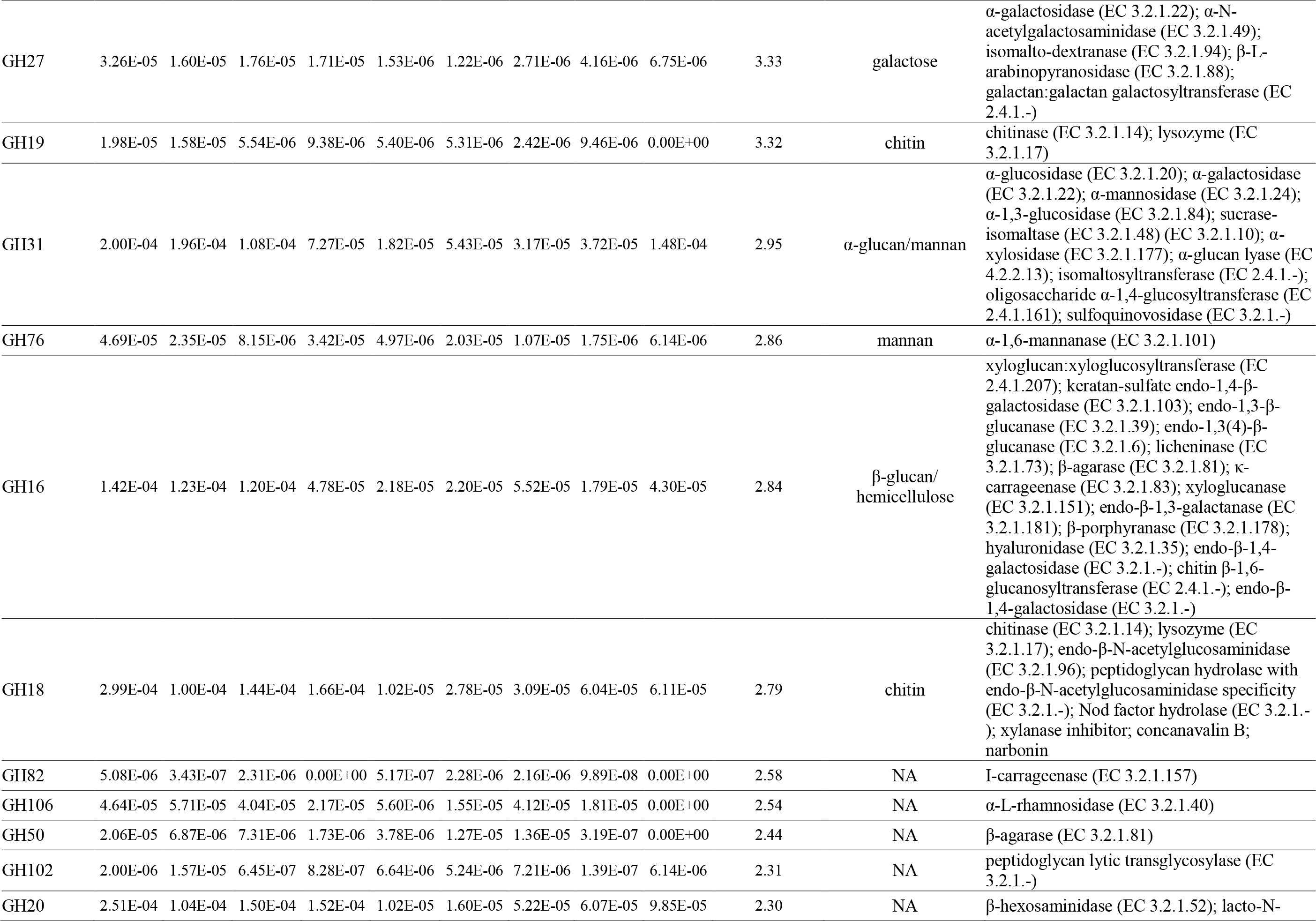

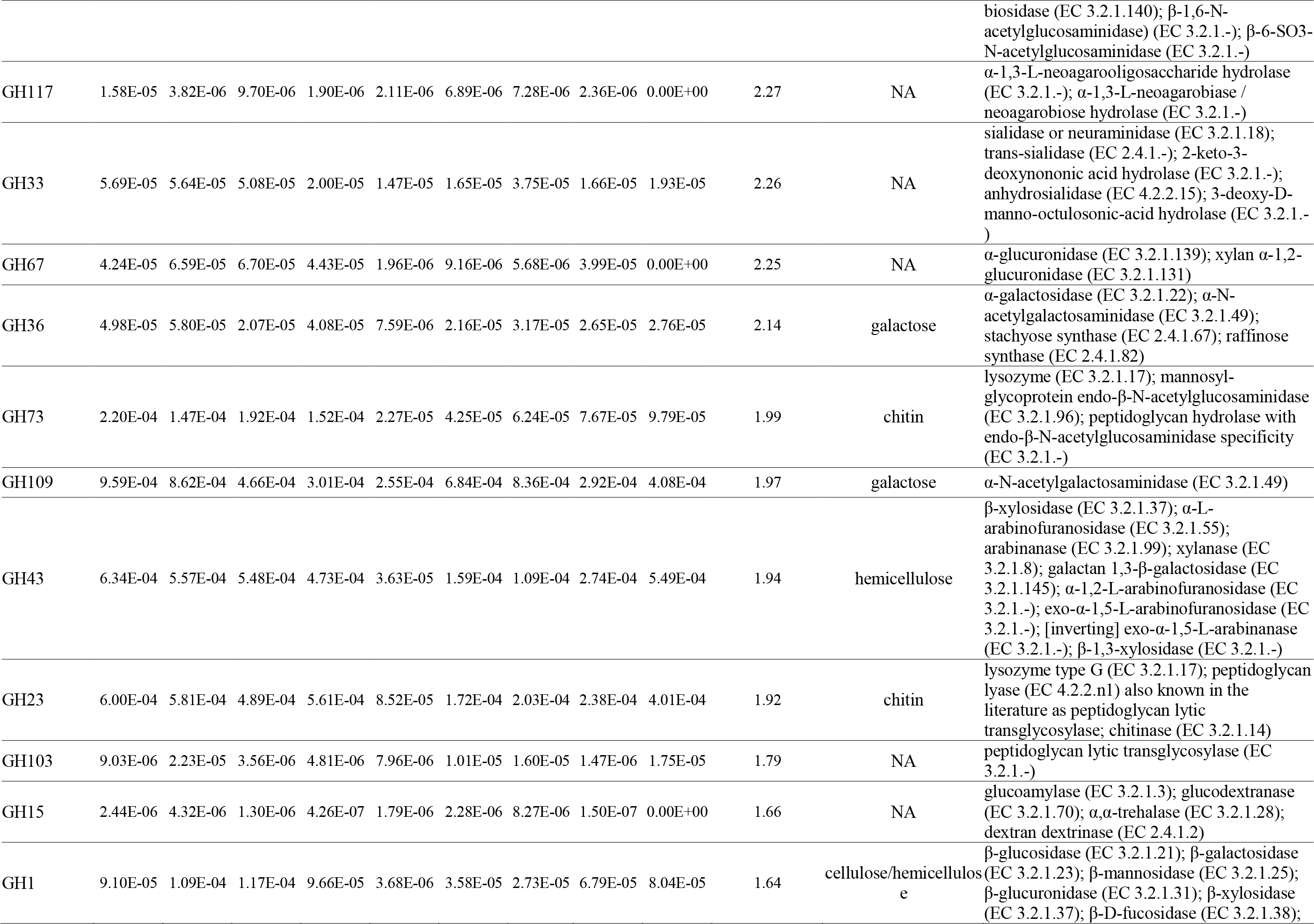

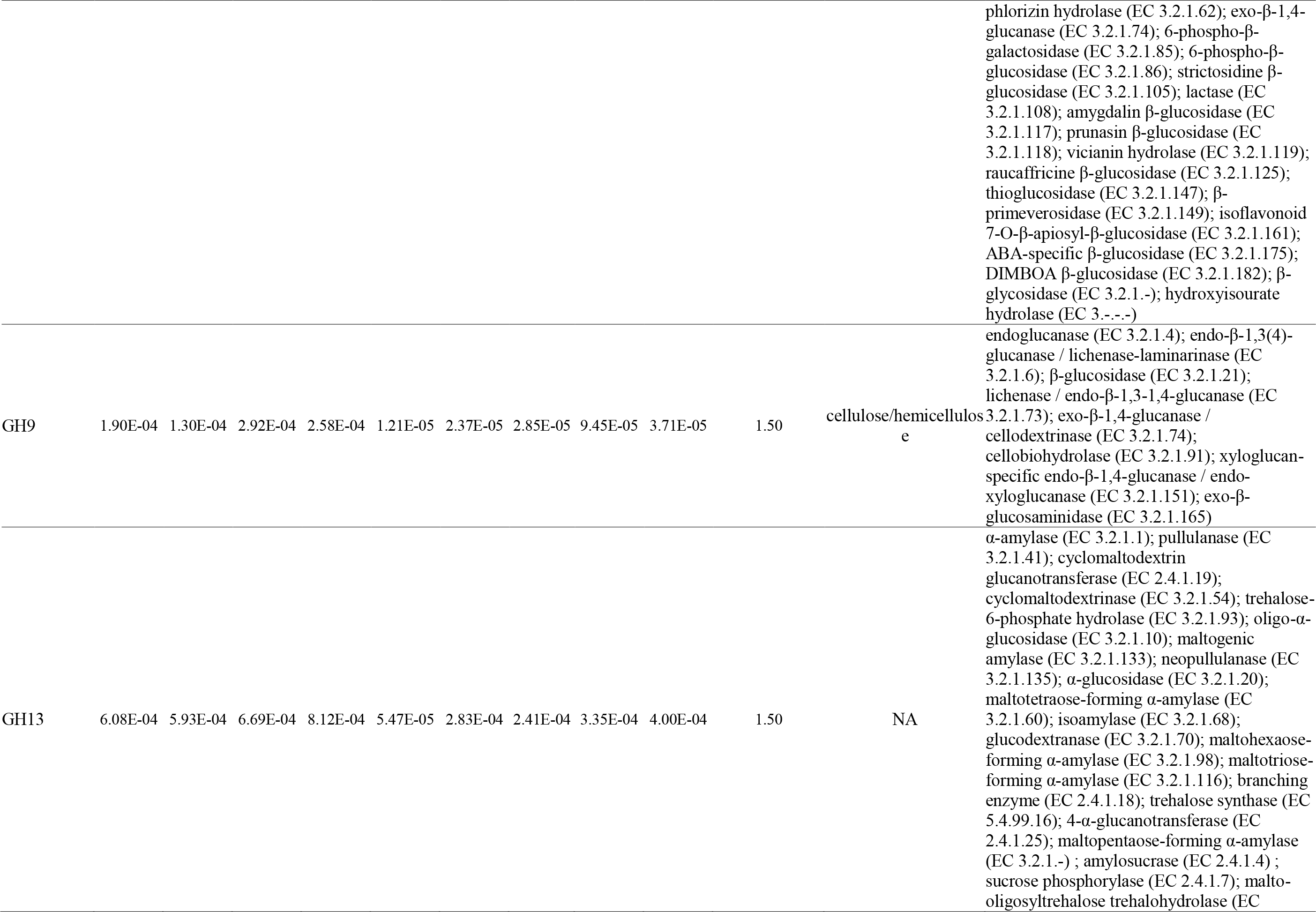

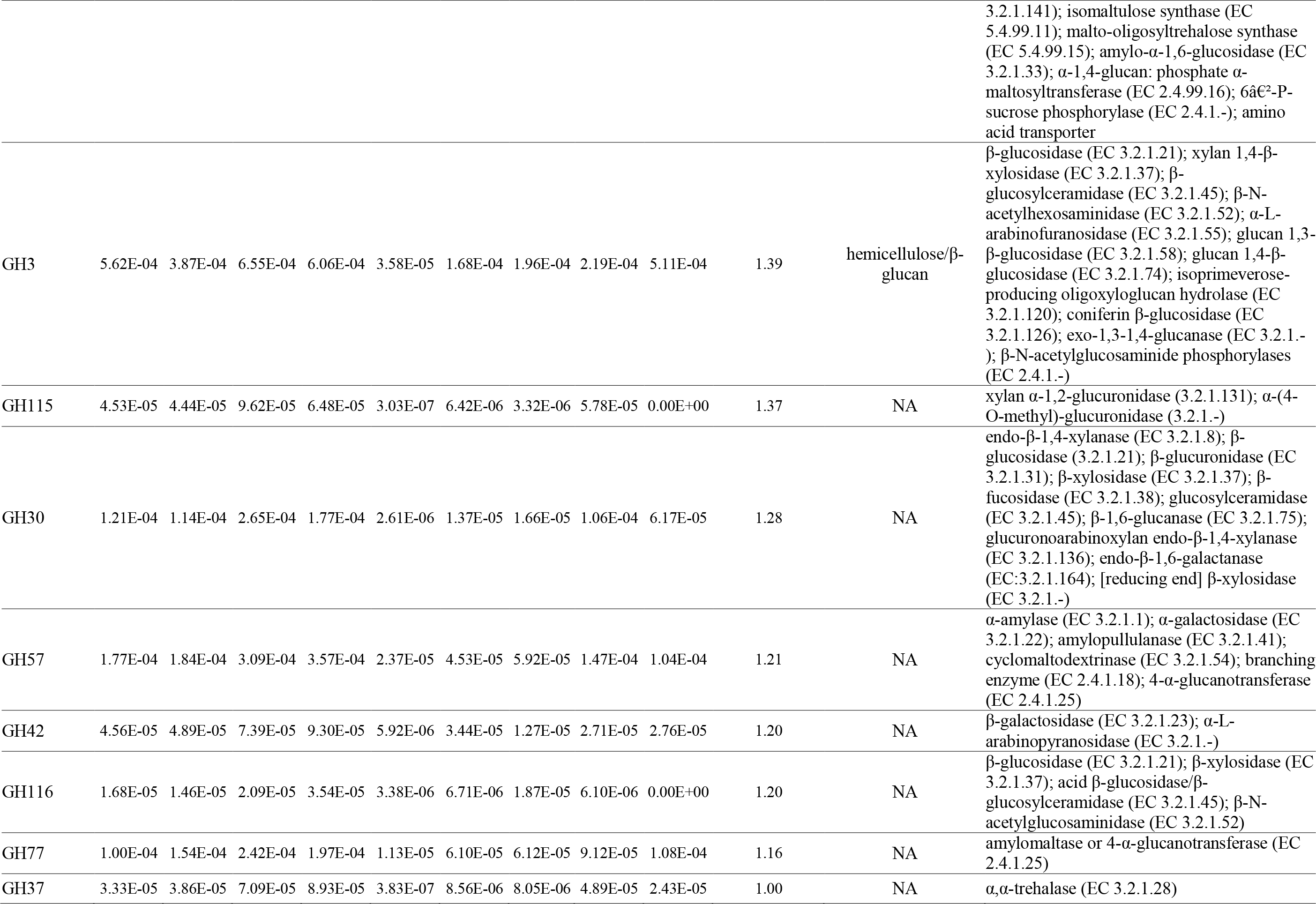

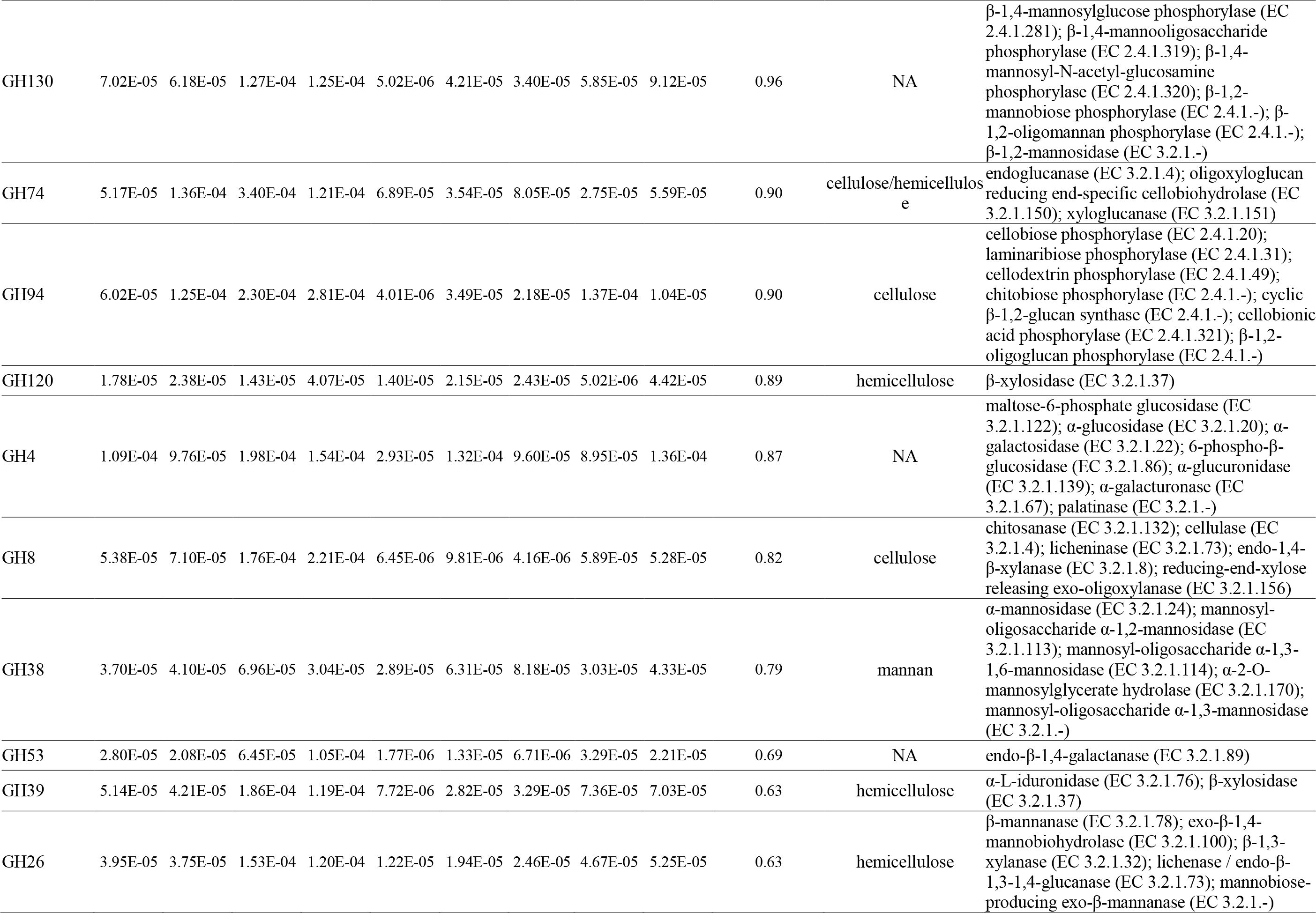

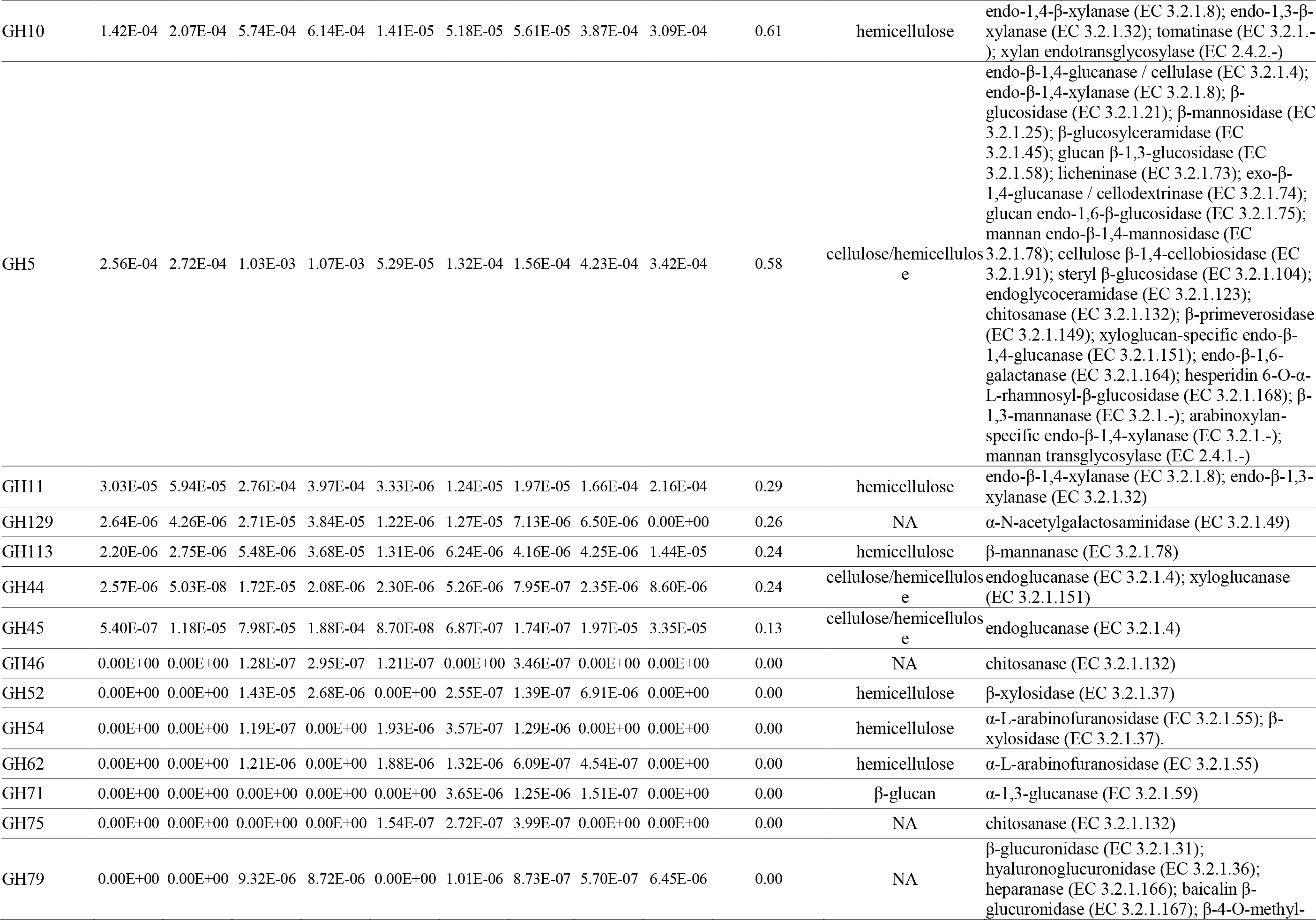

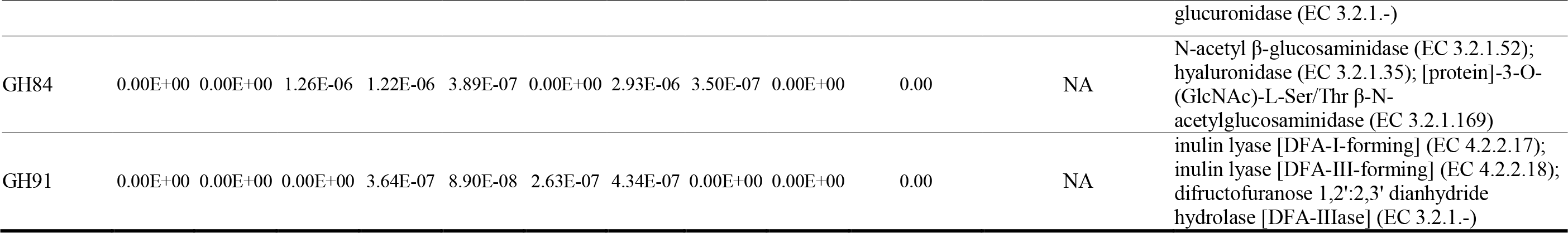

**Table S3.**
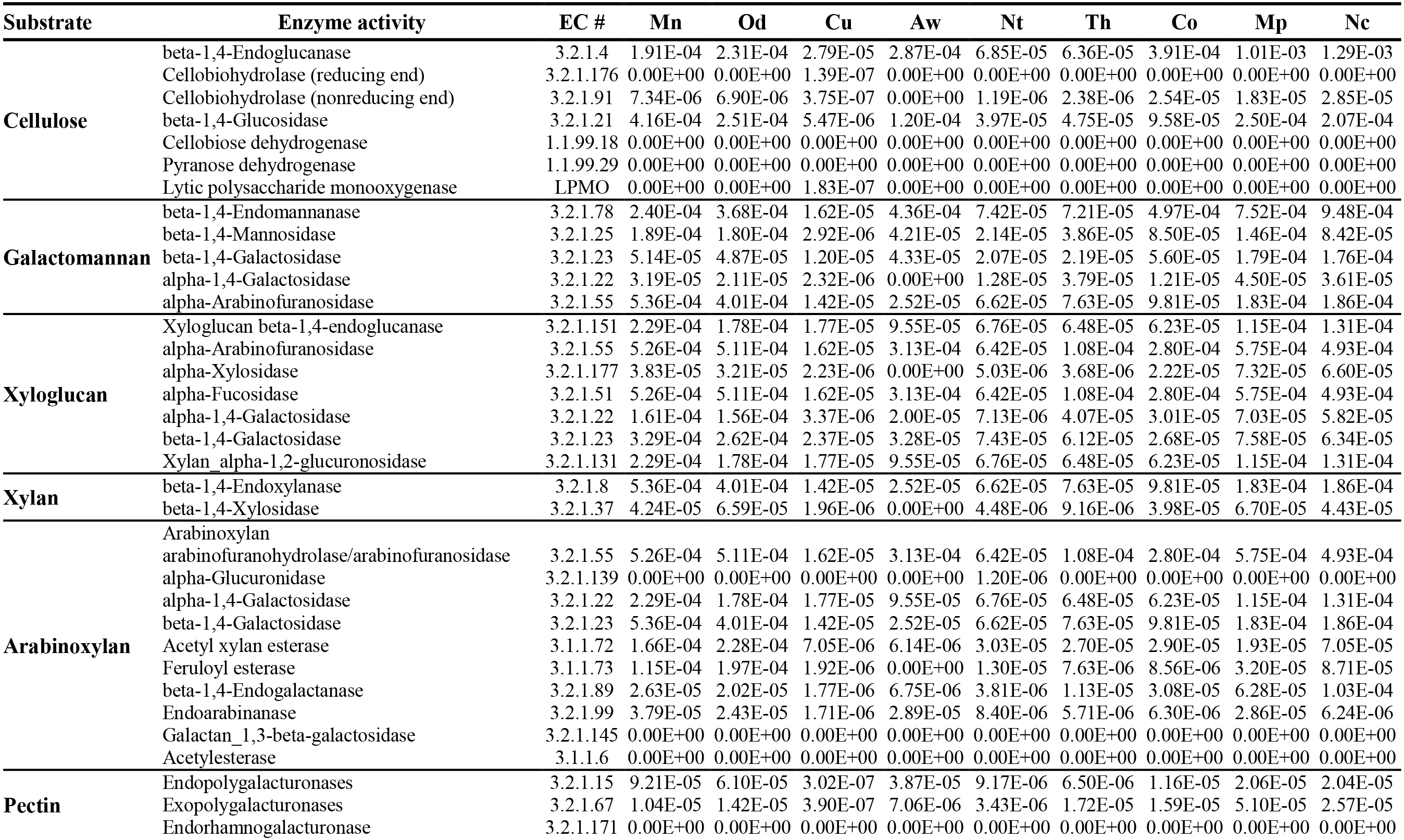
Relative abundance of plant-degrading enzymes in gut metagenomes of nine termites with different diets. Mn: *Macrotermes natalensis*, Od: *Odontotermes* sp., Nc: *Nasutitermes corniger*, Aw: *Amitermes wheeleri*, Mp: *Microcerotermes parvus*, Co: *Cornitermes* sp., Th: *Termes hospes*, Nt: *Neocapritermes taracua*, Cu: *Cubitermes ugandensis*.

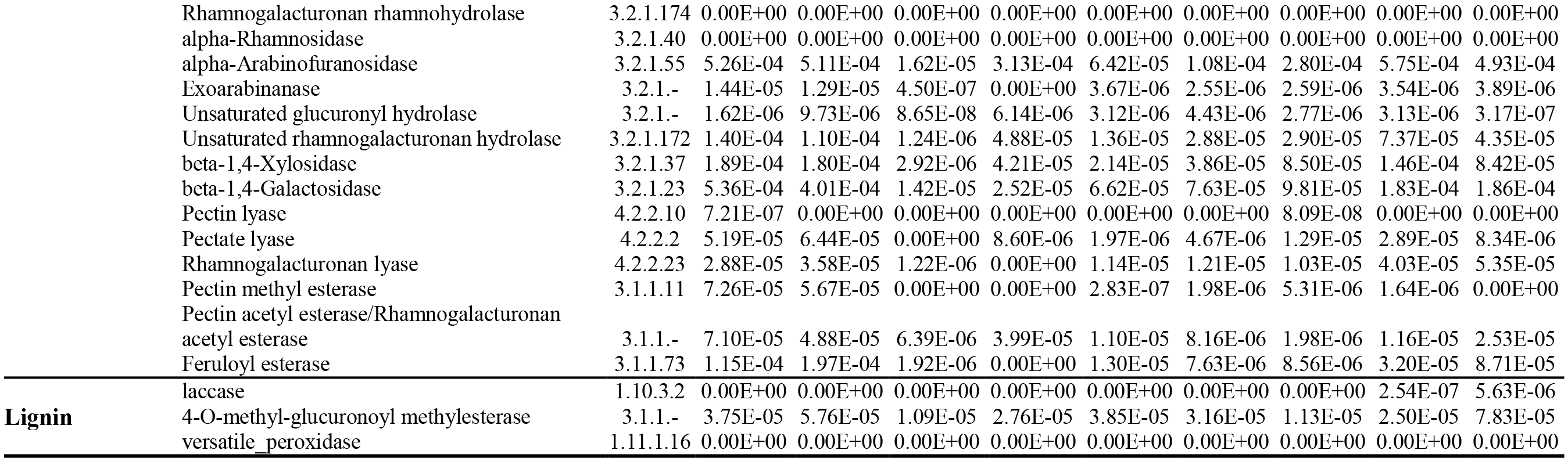

**Table S4.** Relative abundance of the main catalytic types of proteases in the gut metagenomes of nine termite species with different diets. Mn: *Macrotermes natalensis*, Od: *Odontotermes* sp., Nc: *Nasutitermes corniger*, Aw: *Amitermes wheeleri*, Mp: *Microcerotermes parvus*, Co: *Cornitermes* sp., Th: *Termes hospes*, Nt: *Neocapritermes taracua*, Cu: *Cubitermes ugandensis*.

| Type of protease | Mn | Od | Cu | Aw | Nt | Th | Co | Mp | Nc |
| --- | --- | --- | --- | --- | --- | --- | --- | --- | --- |
| Aspartic Peptidases | 5.66E-04 | 5.75E-04 | 4.36E-04 | 3.93E-04 | 3.78E-04 | 3.94E-04 | 2.60E-04 | 4.69E-04 | 4.59E-04 |
| Cysteine Peptidases | 5.03E-03 | 5.29E-03 | 1.29E-03 | 4.86E-03 | 2.35E-03 | 2.74E-03 | 2.81E-03 | 4.33E-03 | 4.16E-03 |
| Glutamic Peptidases | 1.67E-06 | 6.38E-06 | 3.41E-06 | 1.00E-07 | 2.32E-06 | 8.36E-07 | 1.00E-07 | 1.00E-07 | 2.66E-07 |
| Metallo Peptidases | 1.09E-02 | 1.14E-02 | 2.74E-03 | 9.83E-03 | 7.24E-03 | 6.07E-03 | 4.94E-03 | 1.07E-02 | 1.01E-02 |
| Asparagine Peptide Lyases | 8.28E-04 | 6.91E-04 | 7.94E-05 | 4.90E-04 | 3.12E-04 | 3.29E-04 | 2.57E-04 | 5.64E-04 | 5.96E-04 |
| Mixed Peptidases | 2.00E-05 | 2.62E-05 | 6.04E-06 | 4.91E-06 | 1.56E-05 | 8.35E-06 | 2.92E-06 | 9.05E-06 | 1.26E-05 |
| Serine Peptidases | 9.39E-03 | 9.34E-03 | 2.13E-03 | 7.57E-03 | 5.30E-03 | 4.54E-03 | 4.16E-03 | 6.63E-03 | 6.52E-03 |
| Threonine Peptidases | 4.85E-04 | 2.89E-04 | 6.51E-04 | 6.09E-04 | 1.09E-03 | 7.34E-04 | 4.81E-04 | 4.97E-04 | 6.56E-04 |
| Peptidases of Unknown Type | 1.20E-03 | 1.21E-03 | 3.46E-04 | 9.15E-04 | 7.93E-04 | 7.31E-04 | 5.45E-04 | 6.97E-04 | 7.08E-04 |
| Sum | 2.85E-02 | 2.88E-02 | 7.69E-03 | 2.47E-02 | 1.75E-02 | 1.56E-02 | 1.35E-02 | 2.39E-02 | 2.32E-02 |

